# Interdigitated immunoglobulin arrays form the hyperstable surface layer of the extremophilic bacterium *Deinococcus radiodurans*

**DOI:** 10.1101/2022.09.15.508085

**Authors:** Andriko von Kügelgen, Sofie van Dorst, Keitaro Yamashita, Danielle L. Sexton, Elitza I. Tocheva, Garib Murshudov, Vikram Alva, Tanmay A. M. Bharat

**Author notes:** **Correspondence to** Tanmay A.M. Bharat, **Email:**.

## Abstract

*Deinococcus radiodurans* is an atypical diderm bacterium with a remarkable ability to tolerate various environmental stresses, partly because of its complex cell envelope encapsulated within a hyperstable surface layer (S-layer). Despite decades of research into this cell envelope, atomic structural details of the S-layer have remained obscure. In this study, we report the electron cryomicroscopy structure of the *D. radiodurans* S-layer, showing how it is formed by the Hexagonally Packed Intermediate-layer (HPI) protein arranged in a planar hexagonal lattice. The HPI protein forms an array of immunoglobulin-like folds within the S-layer, with each monomer extending into the adjoining hexamer, leading to a highly interconnected, stable, sheet-like arrangement. Using electron cryotomography and subtomogram averaging from focused ion beam-milled *D. radiodurans* cells, we obtained a structure of the cellular S-layer, showing how this HPI S-layer coats native membranes on the surface of cells. Our S-layer structure from the diderm bacterium *D. radiodurans* shows similarities to immunoglobulin-like domain-containing S-layers from monoderm bacteria and archaea, highlighting shared traits in cell surface organization across different domains of life, with connotations on the evolution of immunoglobulin-based molecular recognition systems in eukaryotes.

## Introduction

*Deinococcus radiodurans* is an atypical diderm bacterium with a remarkable ability to endure various environmental stresses, including nuclear radiation, extreme temperatures, vacuum, oxidation, and desiccation (1, 2). In fact, it can tolerate acute doses of ionizing radiation of 5000 Grays (Gy) with no loss of viability (2). By comparison, doses of 5 Gy are fatal for humans and 200-800 Gy for *Escherichia coli* (3). Due to its extreme tolerance to radiation and desiccation, not only can *D. radiodurans* survive under extreme conditions on earth but also in outer space for years (4). Therefore, *D. radiodurans* has been of tremendous interest for biotechnological applications such as bioremediation of radioactive waste. The biochemical mechanisms underlying the hyperstability of *D. radiodurans* have not been fully elucidated yet. However, its efficient mechanisms to repair damaged DNA (5), its molecular machinery to prevent oxidative protein damage (6, 7), and its complex cell envelope are thought to be important factors promoting hyperstability (8–12).

*D. radiodurans* is the prototypical species of the evolutionarily deep-branching genus Deinococcus, which is considered an intermediate in the diderm-monoderm transition, exhibiting characteristics of both Gram-negative and Gram-positive bacteria (2). While *D. radiodurans* stains Gram-positive because it contains a thick peptidoglycan (PG) layer covering the inner membrane, its cell envelope resembles that of Gram-negative bacteria in possessing a periplasmic space and an outer membrane (OM) (8–10). Additionally, the OM of *D. radiodurans* is covered by a carbohydrate-coated, hexagonal surface layer (S-layer) (13), which is a proteinaceous, paracrystalline coat frequently found on the surface of archaea and bacteria (14). In general, S-layers are involved in critical cellular functions such as preservation of cell shape, protection from phages, biomineralization, cell adhesion, and induction of virulence (15). The S-layer in *D. radiodurans* has been reported to be composed entirely of the <u>H</u>exagonally <u>P</u>acked Intermediate-layer (HPI) surface protein and exhibits impressive hyperstability at high temperatures (> 80 °C) even in the presence of strong detergents such as sodium dodecyl sulfate (SDS) and urea (13, 16). While several pioneering studies have revealed the molecular envelope formed by the *D. radiodurans* S-layer using three-dimensional electron microscopy (EM) (9, 11, 13, 17–22), the atomic structure of the S-layer, the principles governing its assembly, and its association with other components of the cell envelope remain elusive.

In this study, we report the cryo-EM structure of the S-layer from *D. radiodurans*. Our structure confirms that this S-layer is solely made up of the HPI protein, arranged in a hexagonal lattice, with no density observed for additional proteins. The HPI protein within the S-layer is arranged as an interdigitated array of immunoglobulin-like (Ig-like) domains, with the inter-hexameric linkages formed by one Ig-like domain swapping into the adjoining hexameric center, leading to a highly interconnected sheath-like arrangement. Cryo-electron tomography (cryo-ET) of focused ion beam (FIB)-milled cells further support our cryo-EM structure and confirm the organization of the HPI proteins within the cellular *D. radiodurans* S-layer. Comparing the S-layer structure from *D. radiodurans* to the ones from Gram-positive bacteria *Bacillus anthracis* (23) and *Geobacillus stearothermophilus* (24, 25) and the archaeon *Haloferax volcanii* (26) reveals that Ig-like-domains are present in all those S-layer proteins (SLPs), representing a common feature in the arrangement of many prokaryotic S-layers. Our observations suggest a common mechanism for cell surface organization shared across different domains of life, with implications for understanding the evolution of eukaryotic molecular recognition systems based on immunoglobulins.

## Results

### Cryo-EM structure of the *D. radiodurans* HPI S-layer

To elucidate the organization of the cell envelope in *D. radiodurans*, we proceeded to solve the atomic structure of the purified S-layer. For this purpose, we isolated native S-layer (Materials and Methods) by adapting a previously described protocol that employed SDS solubilization (8, 13). Images of the isolated S-layer revealed top and side views of S-layer-like two-dimensional sheets displaying a characteristic planar hexagonal symmetry (Figure 1A). We used a previously described single-particle cryo-EM data analysis workflow (26) to resolve a global 2.5 Å-resolution structure of the S-layer within the planar sheets (Figure 1B-D, S1, and Table S1). The 2.5 Å-resolution map allowed us to build an atomic model of the *D. radiodurans* S-layer (Figure 1C-E and Movie S1).

**Figure 1.**
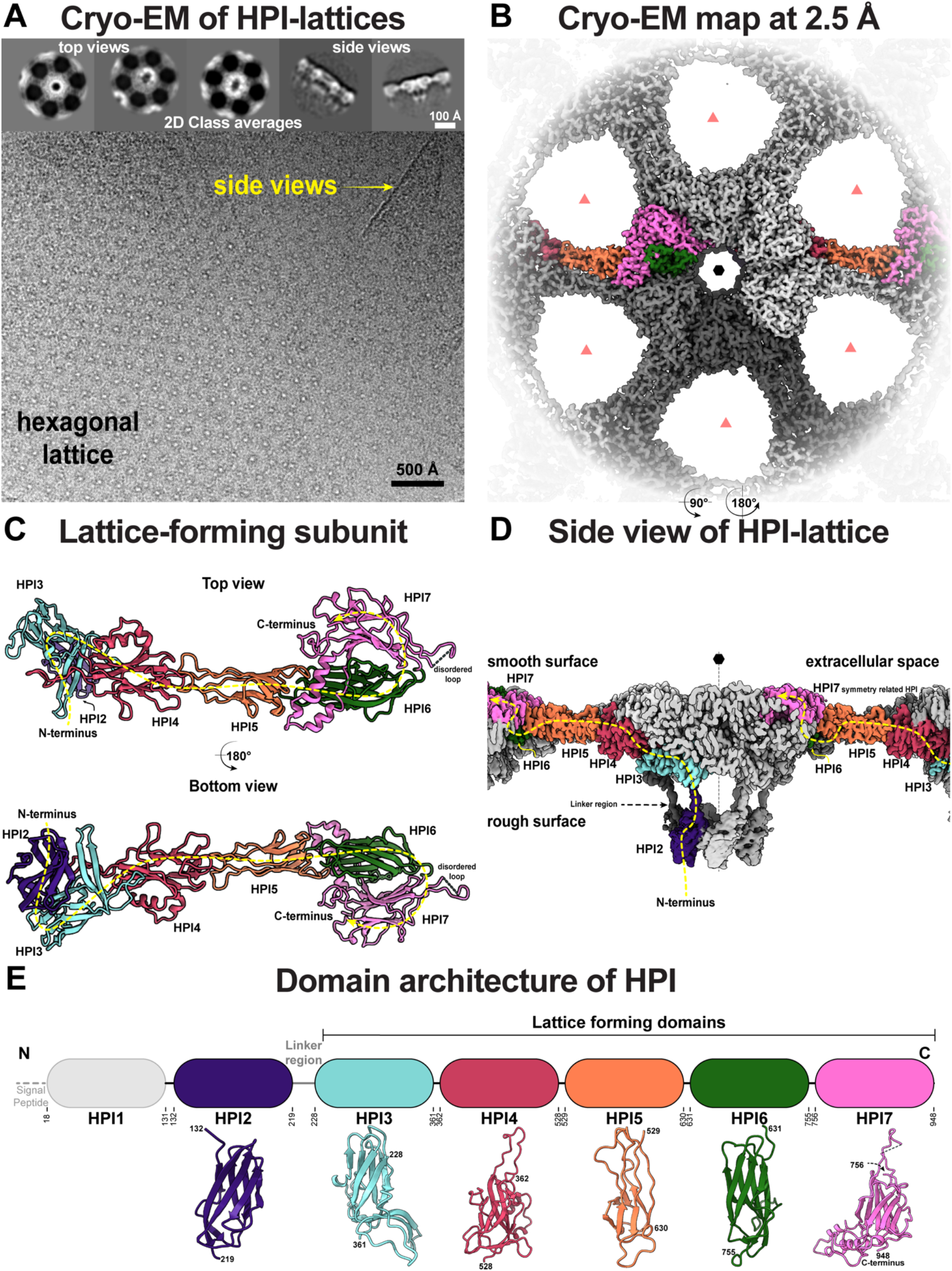
Cryo-EM reconstruction of *D. radiodurans* S-layer. (A) Cryo-EM images of purified *D. radiodurans* cell envelopes show hexagonal sheets of S-layer, in line with previous reports. Insets – characteristic top and side views observed in class averages. (B) Density map of the S-layer is shown from the top, with two (two-fold) symmetry-related copies of SLP monomers shown in full color. The resolution of the map is 2.5 Å (see Figure S1). The nearly hexameric (black hexagon) and trimeric symmetry (light red triangles) axes are marked. During cryo-EM refinement only six-fold symmetry was imposed. (C) The S-layer is made up of repeating copies of a single SLP called HPI, shown as a ribbon diagram. The path of one monomer is shown with a dotted yellow line. (D) The HPI S-layer cryo-EM map shown in an orthogonal orientation compared to panel B, a view along the lattice. (E) Schematic cartoon of the HPI protein sequence together with resolved cryo-EM structures of individual HPI domains. HPI1 is not resolved in our cryo-EM map and is likely involved in anchoring the lattice to the cell.

The atomic model of the S-layer shows that only the HPI protein forms the S-layer, with no additional density observed in the map (Figures 1B, 1D and S1), in line with previous reports (8, 9, 11–13, 20–22). At the sequence level, the HPI protein is composed of seven β-strand-rich immunoglobulin (Ig)-like domains (HPI1 to HPI7 from hereon, Figure 1E and S2), of which six (HPI2-HPI7) were resolved in our cryo-EM map and revealed an extended arrangement of the HPI protein. HPI1 domain (residues 18-132) was not resolved in our structure, suggesting flexibility in its position relative to the S-layer lattice. To obtain structural information on HPI1 and its connectivity to HPI2, we built a structural model of full-length HPI using AlphaFold2 (27). The predicted model shows that HPI1 adopts an Ig-like fold and probably connects to HPI2 through a long, disordered linker (residues 114-131) (Figure S3). The first experimentally resolved domain, HPI2 (residues 132-219), is connected to HPI3 through a long, stretched linker containing residues 220-229 (Figure 1D). HPI3-HPI7 form a planar sheet (Figures 1D, 2, and S1) and are highly variable compared to HPI1 and HPI2 due to extensive insertions (Figure 1E). Although the seven Ig-like domains of HPI are quite sequence divergent to each other, they exhibit similar topologies and resemble the ‘Immunoglobulin/Fibronectin type III/E set domains/PapD-like’ topology group (T-group) in the Evolutionary Classification of Protein Domains (ECOD) database (28).

The domains of HPI show a bipartite arrangement parted by a long linker (residues 220-229, Figure 1D) that separates HPI1-HPI2 from HPI3-HPI7, suggesting that while the latter domains assemble the canopy of the S-layer, the former ones form the OM-anchoring stalk of the S-layer (14). Indeed, HPI is predicted to possess a C-terminal lipoprotein signal peptide with a canonical four-residue-long lipobox motif and a conserved cysteine residue. Lipobox-containing lipoproteins are widespread in bacteria and have been previously characterized to be anchored to the OM by a post-translational lipid modification of the conserved cysteine residue, which forms the N-terminus of the mature protein after cleavage of the signal peptide (29). In fact, in the mature HPI protein, the conserved cysteine constitutes the first residue, implying that HPI might be anchored to the OM by lipidation (Figure S2).

### HPI monomers from an interdigitated array in the *D. radiodurans* S-layer

The extended monomers of HPI are packed in a sheet as an interdigitated array (Figure 2A-C and Movie S1). Each monomer spans two hexamers of the S-layer lattice via HPI5, which is arranged as a two-fold symmetric bridge linking adjoining hexamers (Figure 2C). This extended arrangement partly explains the exceptional stability of the *D. radiodurans* S-layer, since there are large protein:protein interfaces making up the lattice (Figure 2C). Each hexamer of the S-layer lattice itself is made up of two stacked tiers, the first formed by HPI3 and HPI4, connected by the HPI5 dimeric bridge, and the second (top) formed by HPI6 and HPI7 (Figures 2D and S4A).

**Figure 2.**
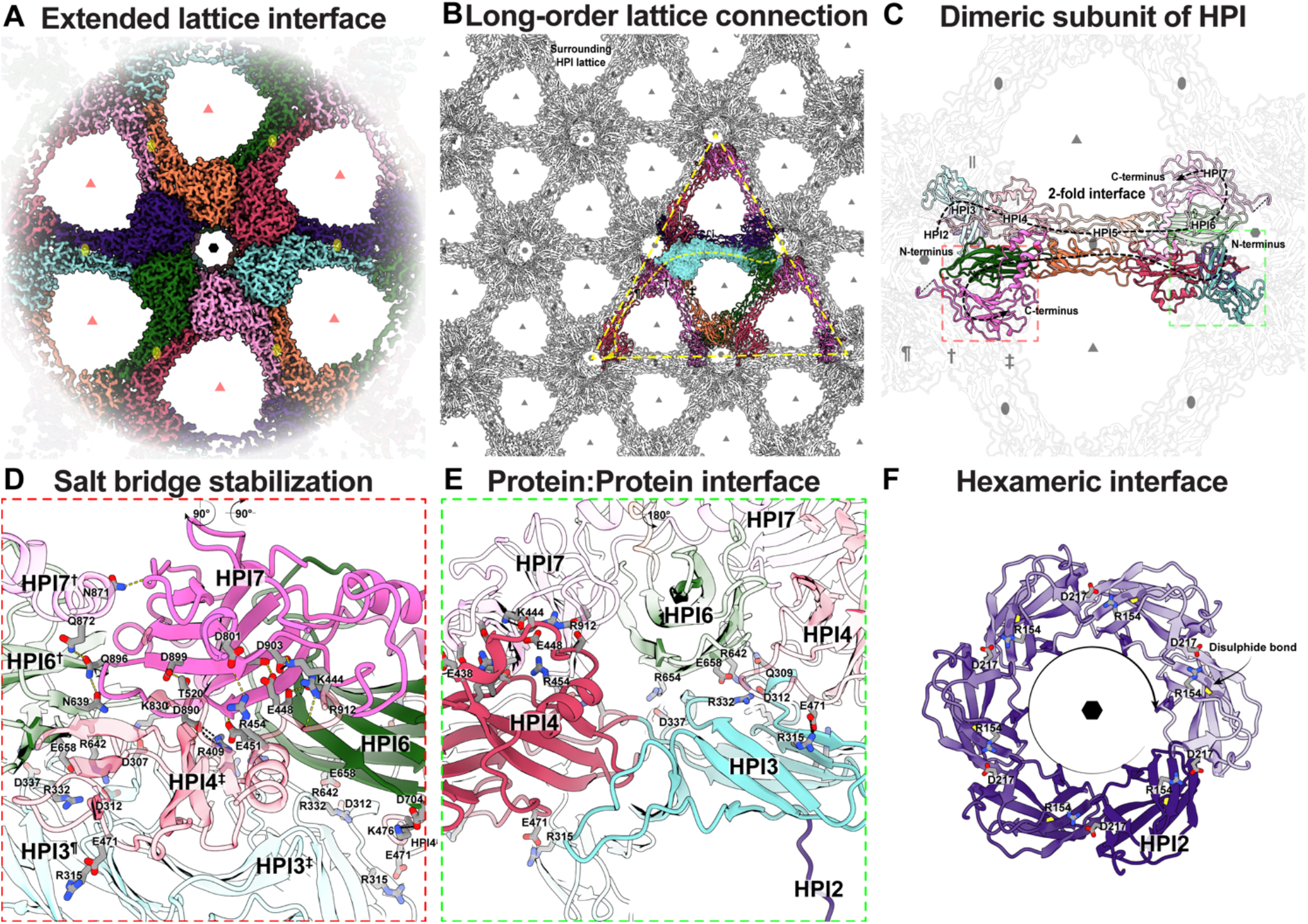
HPI Ig-like domains form a highly interconnected lattice stabilized by multiple salt bridges. (A) Cryo-EM density of an HPI hexamer. Different HPI monomers are colored differently with the hexameric, trimeric, and dimeric axes marked by a black hexagon, light red triangles, and yellow ovals, respectively. (B) Each HPI monomer (cyan) interacts tightly with nine other monomers, spanning six hexamers of the lattice, leading to a highly interconnected S-layer. (C) Each HPI monomer swaps into the adjoining hexamer through the extended HPI5 domain. The lattice is stabilized by multiple salt bridges, shown in enlarged views marked with a (D) red box, and (E) green box. (F) The hexameric HPI2 domain is also stabilized by several salt bridges.

This extended arrangement of HPI monomers in the lattice results in a porous lattice with large gaps (Figure 2A-B). The hexameric pore is relatively large (∼33 Å) compared to the S-layer structure reported for the diderm Gram-negative *C. crescentus* (20 Å, Figure S5A-B). There are additional gaps between the *D. radiodurans* S-layer hexamers that were even larger (∼76 Å), resulting in a relatively holey structure, suggesting that this S-layer is not acting to occlude small molecules from the cell surface (Figure 2A-B). In the Gram-negative S-layer lattice of *C. crescentus*, the hexameric, trimeric, and dimeric interfaces making up the lattice are lined with Ca^2+^ ions, which have been shown to be essential for lattice assembly and stabilization (30–32). In the same vein, some putative cation-binding sites are also seen in the HPI lattice (Figure S5C-D), which may play a role in stabilizing the lattice. Additionally, the S-layer is held strongly by many salt bridges forming an extensive network through the HPI lattice (Figures 2D-F and S4B). This network of salt bridges likely plays a key role in the observed hyperstability of the *D. radiodurans* S-layer. Finally, there are two cysteine residues in close proximity (C554 and C666) between HPI5 and HPI6 of each monomer (Figure S5E). Although densities connecting these residues were not observed in the map, they could form a putative disulfide bridge joining HPI5 to HPI6, further interconnecting the lattice (Figure 2D-F).

### Ig-like arrays are observed in S-layers across bacterial and archaeal phyla

To investigate whether HPI-like S-layers are found in other bacteria, especially of the Deinococcus-Thermus phylum, we used the Basic Local Alignment Search Tool (BLAST) to search for homologs of HPI in the nonredundant (nr) protein sequence database at NCBI. The search found about 20 homologs, primarily from bacteria of the order Deinococcales, including *Deinococcus wulumuqiensis, Deinoccus phoenicis, Deinoccus murrayi, Deinococcus fonticola*, as well as *Deinobacterium chartae*, which possesses two paralogs (Figures 3 and S2 and Table S2). The search also identified a homolog in *Thermus thermophilus* (NCBI ID BDG20071.1), but it is unknown whether it forms an S-layer *in vivo*. The sparse occurrence of HPI indicates that HPI-like S-layers may not be widespread in the Deinococcus-Thermus phylum or that the sequences of many HPI proteins may have diverged significantly, making their detection difficult.

**Figure 3.**
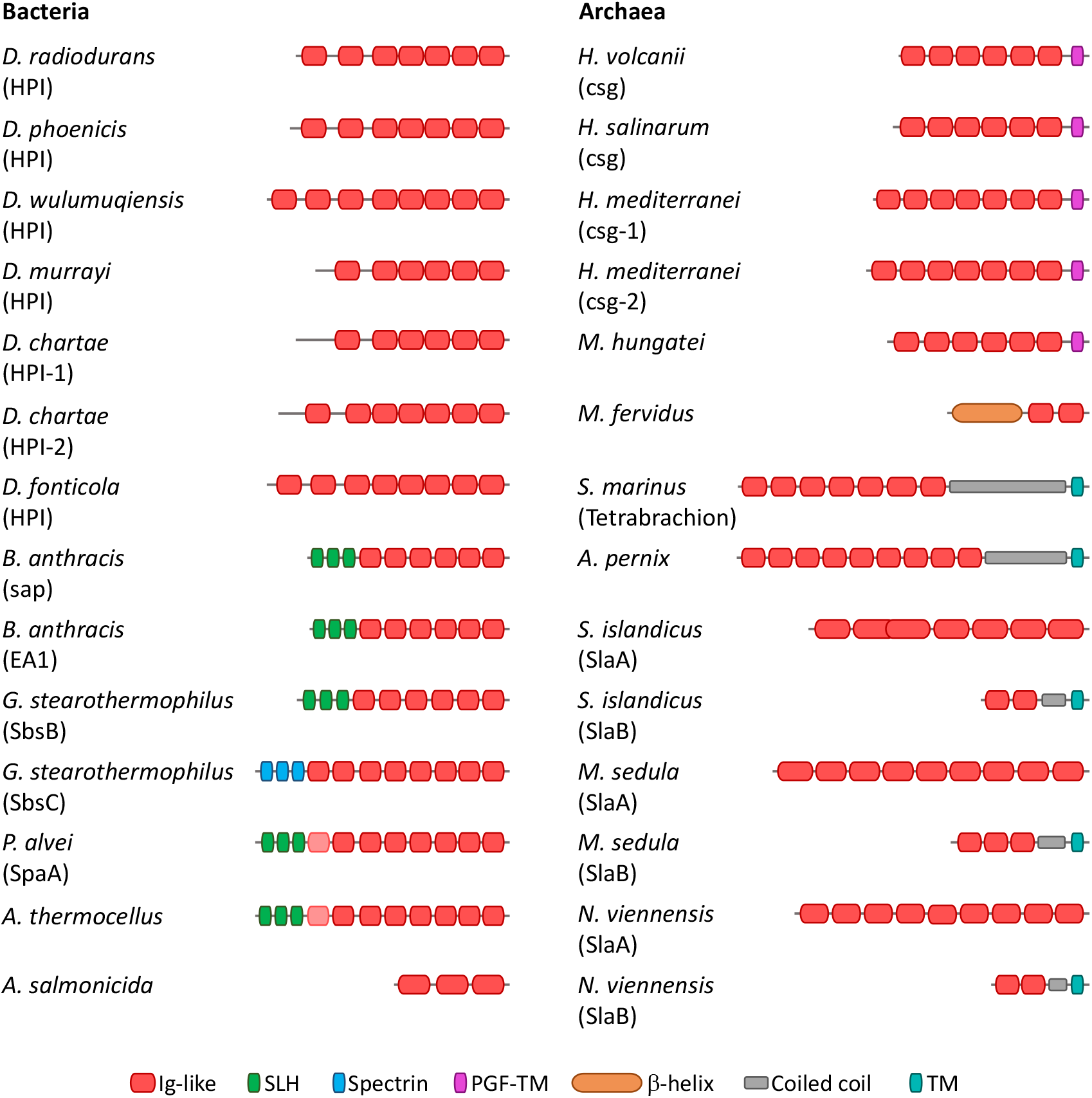
Domain organization of prokaryotic Ig-like domain-containing SLPs. The Ig-like domain arrays observed in the *D. radiodurans* HPI protein are found across different bacterial and archaeal phyla. Although bacterial and archaeal SLPs are similar in containing Ig-like domain, they employ different anchoring mechanisms; for instance, while many Gram-positive bacterial SLPs possess N-terminal PG-binding SLH domains, many archaeal SLPs possess a C-terminal transmembrane helix. In addition to containing Ig-like and anchoring domains, some archaeal SLPs also contain other domains, such as β-helical and coiled-coil segments. The first Ig-like domain of *P. alvei* and *A. thermocellus* SLPs are highly divergent and are therefore colored in a lighter shade of red. Accession details for the shown proteins are provided in Table S2.

The deinococcal HPI proteins identified in the search are fairly conserved and share pairwise sequence identities of 25%-88% with *D. radiodurans* HPI. Like *D. radiodurans* HPI, most homologs contain seven consecutive Ig-like domains, with some possessing additional N- or C-terminal Ig-like domains or lacking an N-terminal domain (Figure 3). For instance, compared to *D. radiodurans*, HPI of *D. murrayi* lacks the first Ig-like domain, whereas HPI of *D. wulumuqiensis* possesses an additional N-terminal Ig-like domain, indicating that the length of the stalk region connecting the OM to the canopy of the S-layer could vary (Figure 3). On the other hand, HPI proteins from *D. fonticola* and *D. chartae* contain an additional Ig-like domain at their C-termini. Furthermore, all HPI homologs are predicted to contain a lipoprotein signal peptide and an invariant N-terminal cysteine residue in the mature protein, strongly suggesting that the protein is anchored to the OM through lipidation. Lastly, the salt bridges and the putative disulfide bridge between HPI5 and HPI6 mentioned above, and a disulfide bridge in HPI2 are highly conserved in all homologs (Figure S2), indicating that all HPI S-layers are probably hyperstable, in the same manner as the *D. radiodurans* HPI S-layer.

In addition to HPI, arrays of consecutive Ig-like domains have also been structurally characterized in the S-layer proteins of the archaeon *H. volcanii* (csg; PDB 7PTR) (26) and the Gram-positive bacteria *B. anthracis* (Sap; PDB 6QX4) and *G. stearothermophilus* (SbsB; PDB 4AQ1 and SbsC; PDB 4UIC) (23, 24). The Ig-like domains of these proteins assume a fold similar to that of the Ig-like domains of HPI and have been classified into the aforementioned T-group ‘Immunoglobulin/Fibronectin type III/E set domains/PapD-like’ in the ECOD database (Figure S3). Additionally, we analyzed the domain composition of many experimentally characterized SLPs employing sensitive sequence searches and structural modeling using AlphaFold2 and found that Ig-like domains are widespread in prokaryotic SLPs (Materials and Methods and Figure 3). These SLPs frequently exhibit different membrane-anchoring mechanisms and occasionally possess additional non-Ig-like domains. For example, many Gram-positive bacterial SLPs contain N-terminal PG-binding SLH domains, whereas archaeal SLPs are anchored by lipidation of their C-terminal end or through a C-terminal transmembrane helix (14). Finally, although some prokaryotic SLPs without Ig-like domains have also been structurally characterized, for instance, SLPs of the archaeon *Methanosarcina acetivorans* (33), the Gram-positive *Clostridioides difficile* (34), and the Gram-negative bacterium *Caulobacter crescentus* (30–32), Ig-like arrays appear to represent the most prevalent protein arrangement found in SLPs of prokaryotes.

### Structure of the S-layer solved directly from cells, *in situ*

To verify that our structure from purified S-layers is consistent with the native assembly of the HPI proteins, we collected cryotomograms of whole cells. Since *D. radiodurans* cells are ∼4 µm in diameter, we first vitrified them on Finder grids and then generated 200-nm thick lamellae using cryo-FIB milling (10). Cryotomograms revealed that the *in situ* appearance of the S-layer is identical to the purified S-layer (Figure 4A), indicating that the native S-layer has the same arrangement as our purified sample.

**Figure 4.**
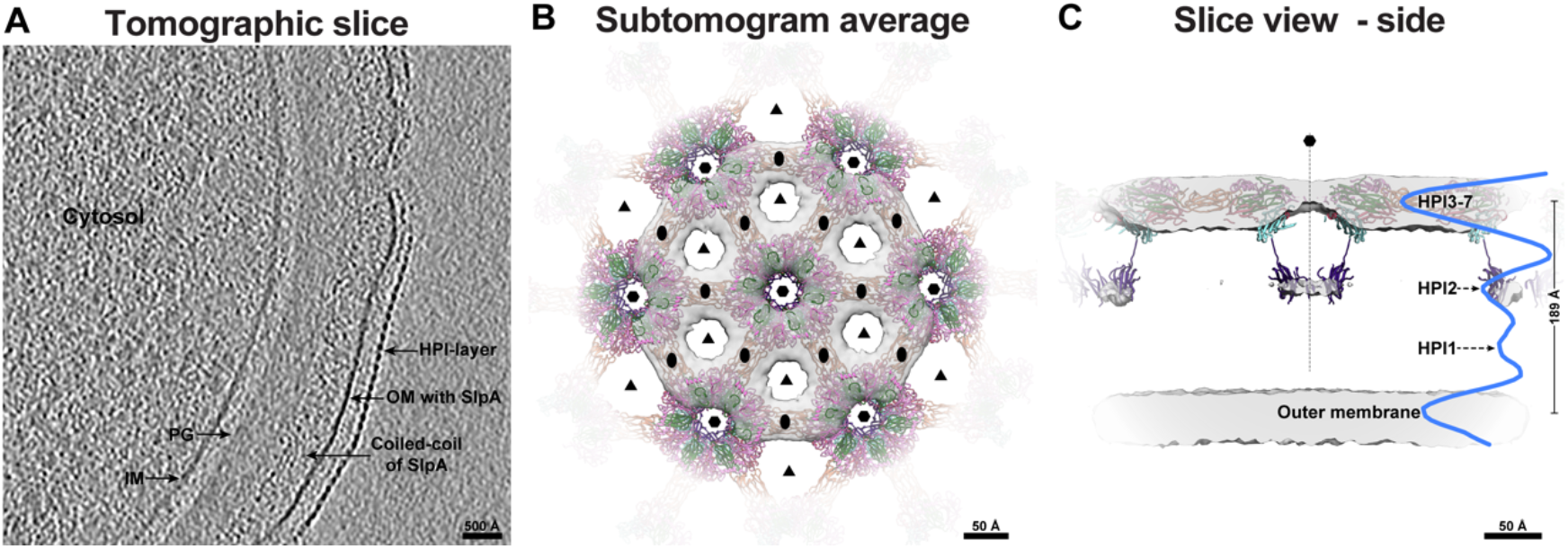
Imaging FIB-milled *D. radiodurans* cells confirms *in vitro* structural data. (A) A tomographic slice through a FIB-milled *D. radiodurans* cell (10) shows the expected density layers on the cell surface corresponding to the S-layer, OM, PG, and IM. (B) Subtomogram averaging of the S-layer confirms the hexagonal arrangement of the S-layer. Docking of the atomic model (ribbon) into the map (grey isosurface) further shows that HPI is the only protein in the S-layer. (C) An orthogonal view of the S-layer subtomogram average shows weak density for HPI1; the density profile is shown in blue.

Next, we used subtomogram averaging of the S-layer lattice using previously described methods (32, 35, 36), producing a 26 Å-resolution reconstruction of the cellular S-layer (Figure 4B-C). Although limited in resolution, our subtomogram averaging agrees with the atomic structure of the HPI S-layer, showing that both are organized in a hexagonal array with the same lattice spacing (Figure 4B-C). The S-layer is seen as large patches coating the cell surface, punctuated with discontinuities, as reported previously (10, 37). The multilayered arrangement of the S-layer was also confirmed by our subtomogram average, with densities for both the canopy (HPI3-7) and the base (HPI2) of the S-layer observed on cells (Figure 4C). Although not resolved in our map, an averaged density profile normal to the OM shows a small peak that may correspond to HPI1 (Figure 4C), where the S-layer is attached to the OM. Also, both the large gaps between hexamers (∼76 Å) and small gaps at the hexameric pore (∼33 Å) were observed in our subtomogram average, which concurs with our cryo-EM atomic model.

Cryotomograms further confirmed that HPI2 is positioned proximal to the cell membrane, and together with HPI1 (not resolved in our average, but seen in the density profile), plays a role in S-layer anchoring. HPI3-HPI7 form the S-layer lattice (the canopy), with large pores exemplifying the lattice. Our *in situ* cellular S-layer structure also confirmed that HPI is the sole protein making up the S-layer, as no density for additional proteins is observed either in our *in vitro* or *in situ* structures.

### Updated model of the *D. radiodurans* cell surface

Using our atomic and cellular data, we report an updated model of the *D. radiodurans* cell surface (Figure 5). Our data are consistent with previous models of the *D. radiodurans* envelope produced primarily by negative-stain EM (8, 9, 11–13, 20–22). The analysis of cryo-ET data of FIB-milled cells further show that HPI is the sole protein making up the S-layer. We show that no densities are present for the highly abundant OM SlpA protein (37), which belongs to the OmpM superfamily of PG-OM tethering proteins (38), in line with previous reports (8, 9, 11–13, 20–22). Further, our data from FIB-milled cells indicates that the S-layer is positioned ∼18 nm away from the OM, in close agreement with recent cryo-EM studies (10, 37). Our atomic structure of the S-layer and the fit of this structure into the subtomogram averaging map of the cellular S-layer show that this lattice coats *D. radiodurans* cells as a highly interconnected sheet with exceptional stability (Figure 5). This updated model of the *D. radiodurans* cell envelope thus allows previous EM studies to be placed into the context of atomic structures.

**Figure 5.**
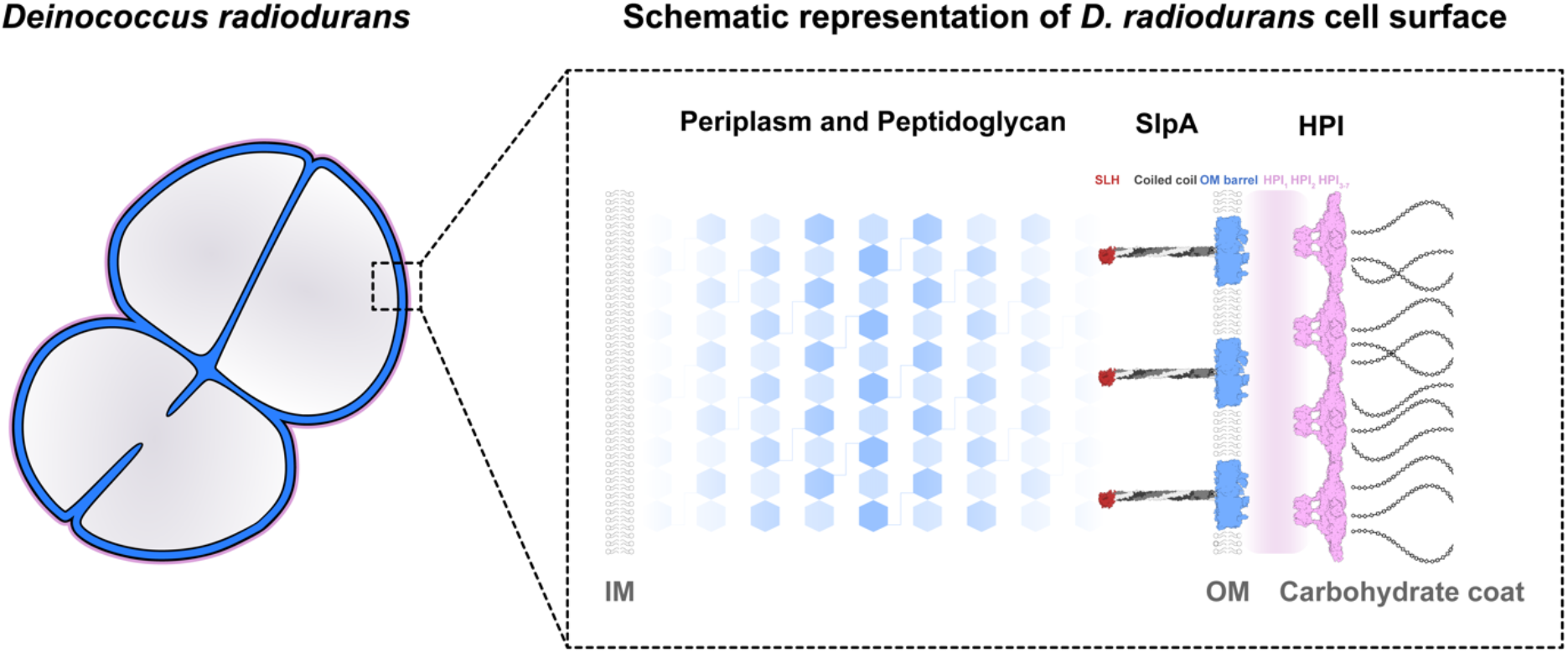
Model of the *D. radiodurans* cell envelope. Schematic model of the *D. radiodurans* cell envelope shows how HPI protein forms the S-layer on cells, in line with previous reports (8, 9, 11–13, 18–22). HPI3-7 form the S-layer lattice farthest away from the cell, while HPI1-2 are closer to and probably tethered to the OM. An abundant protein called SlpA protein is buried in the OM, connecting it to the PG layer via long coiled coils and an N-terminal SLH domain, as shown previously (37, 39). Our results place previous studies into context, and our updated model of the *D. radiodurans* cell envelope will serve as a framework for understanding the cell surface organization of phylogenetically deep-branching bacteria.

## Discussion

The cell envelope of *D. radiodurans* has been studied for over five decades because of its hyperstability and uniqueness compared to other bacteria (17, 22, 37, 39). Numerous studies have employed a variety of approaches to obtain structural and molecular information on the *D. radiodurans* S-layer, its composition (13, 20), symmetry (9, 17, 22), and overall arrangement (11, 12). Here, we build on this knowledge by applying the latest technological advancements in structural biology (40, 41) to obtain an atomic model of the native arrangement of the HPI S-layer.

Our structure places several past studies into context, confirming that the *D. radiodurans* S-layer is made solely of the HPI protein. A recent model of the *D. radiodurans* S-layer proposes that the S-layer incorporates another protein called SlpA (42–44). This, however, is incompatible with our data and also with previous genetic (39), biochemical (45), and EM studies (8, 9, 11–13, 20–22), which have shown that SlpA is a PG-OM tether (39, 45) and that the S-layer is exclusively made up of the HPI protein (8, 9, 11–13, 20–22). Our data support these past studies in that SlpA stabilizes the cell envelope by linking the OM to the PG (37). The proposal that HPI is the sole S-layer protein is verified by our atomic structure (Figures 1–2) and by our *in situ* cryo-ET and subtomogram averaging of native cells (Figure 4), which showed no density corresponding to SlpA in the S-layer.

The HPI protein consists of seven consecutive, evolutionarily divergent Ig-like domains. This organization has been previously observed in Gram-positive bacterial (23, 24) and archaeal (26) SLPs, suggesting a unique ability of Ig-like domains to support diverse functions at the cell surface. It is tempting to speculate that these Ig-like domains may have facilitated the evolution of more complex surface recognition modules (46), such as the eukaryotic immune system. Indeed, it is known that glycosylation patterns on archaeal SLPs help organisms recognize ‘self’ from ‘non-self’ (47). Primordial S-layers built of such Ig domain-containing SLPs may have played important roles in cellular recognition, which in turn could have supported the evolution of the modern eukaryotic cell, as suggested previously (46).

The structure of the *D. radiodurans* S-layer also reveals how this sheet-like assembly displays extreme hyperstability. The domain-swapped arrangement of the hexagonal HPI lattice, in which a single polypeptide chain is shared with adjoining hexamers, leads to a highly interconnected structure that would require multiple biochemical steps for complete disassembly into monomers. Additionally, the lattice is held together by many salt bridges, which provide exceptional stability even in the presence of strong denaturing detergents such as SDS (8, 13). How the S-layer is bound to the OM is unclear, since residues 18-132 of the mature HPI protein were not resolved in our cryo-EM map. However, an invariant cysteine residue near the N-terminus of the mature HPI protein might be lipidated based on our bioinformatic analysis, which agrees with previous reports finding that S-layer preparations stain positive for lipids (9).

This hyperstable *D. radiodurans* S-layer structure could be exploited for synthetic biology applications, allowing surface display of molecules at high-copy numbers on cells and *in vitro*, as previously described for other S-layers (30, 48). This structure also provides important insights into the organization of the enigmatic cell envelope of the phylogenetically deep-branching bacteria and will help illuminate our understanding of how Gram-negative and Gram-positive bacteria evolved by forming the basis for future studies on the arrangement and evolution of prokaryotic cell surfaces.

## Acknowledgments

This work was supported by the Medical Research Council, as part of United Kingdom Research and Innovation (also known as UK Research and Innovation) [Programme MC_UP_1201/31]. For the purpose of open access, the MRC Laboratory of Molecular Biology has applied a CC BY public copyright licence to any Author Accepted Manuscript version arising. T.A.M.B. would like to thank the Human Frontier Science Program (Grant RGY0074/2021), the Vallee Research Foundation, the European Molecular Biology Organization, the Leverhulme Trust, and the Lister Institute for Preventative Medicine for support. E.I.T. was supported by a Natural Sciences and Engineering Research Council of Canada Discovery Grant (RGPIN 04345) and D.L.S. was supported by NSERC Postdoctoral Fellowship (546024). V.A. would like to thank Andrei Lupas for continued support and the Human Frontier Science Program (Grant RGY0074/2021). This work was partly supported by institutional funds of the Max Planck Society.

## Supplementary Figures

**Figure S1.**
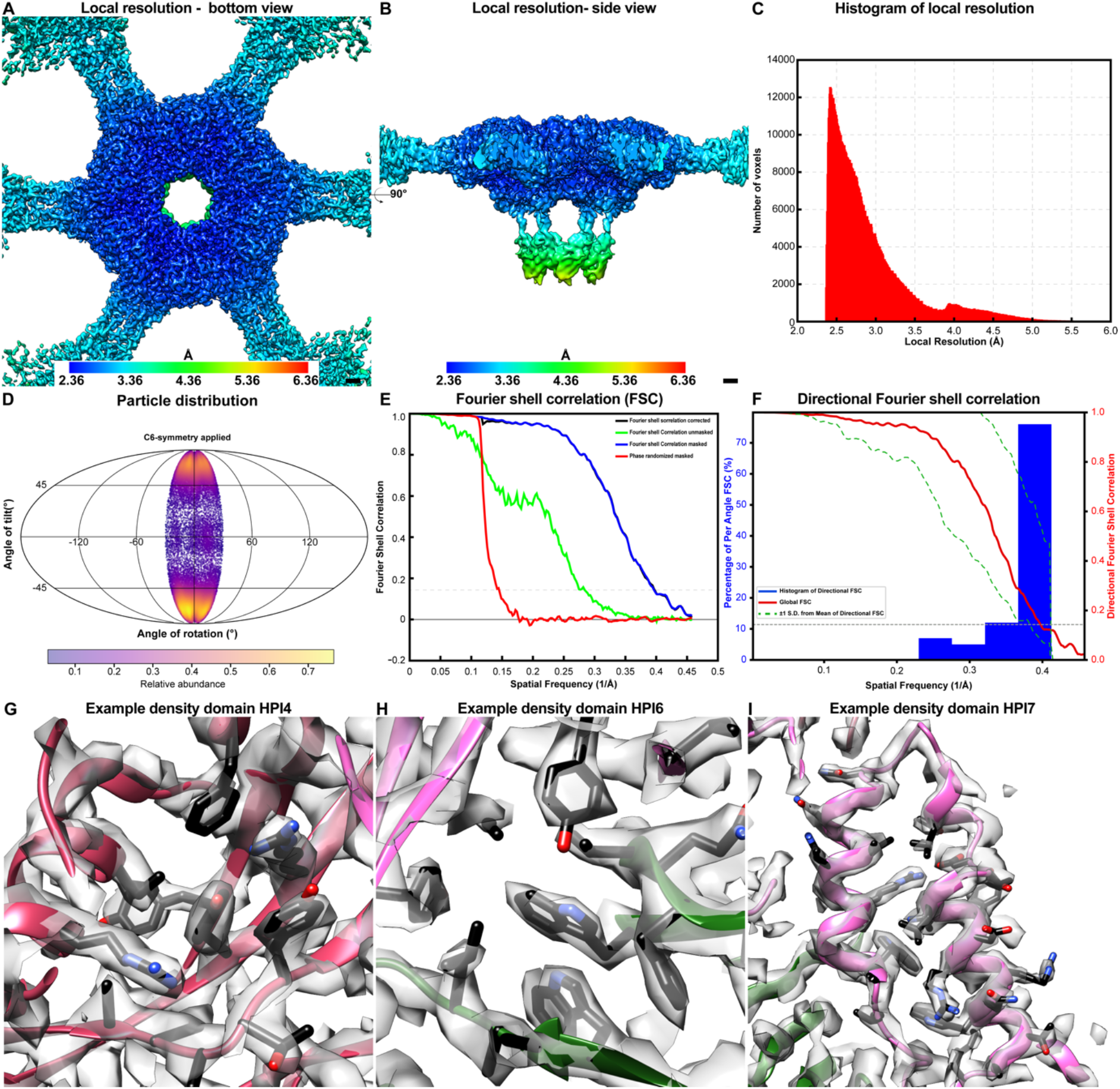
Cryo-EM structure of the *D. radiodurans* S-layer. (A-B) Local resolution of the cryo-EM map estimated in RELION, plotted into the density, shown in two orthogonal orientations. The resolution of HPI2 is slightly lower. (C) Histogram of local resolutions in voxels of the cryo-EM map. (D) Angular distribution of the particles in the data set, shown on a relative scale (purple denotes low and yellow denotes high). (E) Fourier Shell Correlation (FSC) estimation of the resolution of the map. (F) 3D FSC between two random halves of the data (49). (G-I) Examples of the cryo-EM density and the fit of the model into the map. Scale bars: A-B) 10 Å.

**Figure S2.**
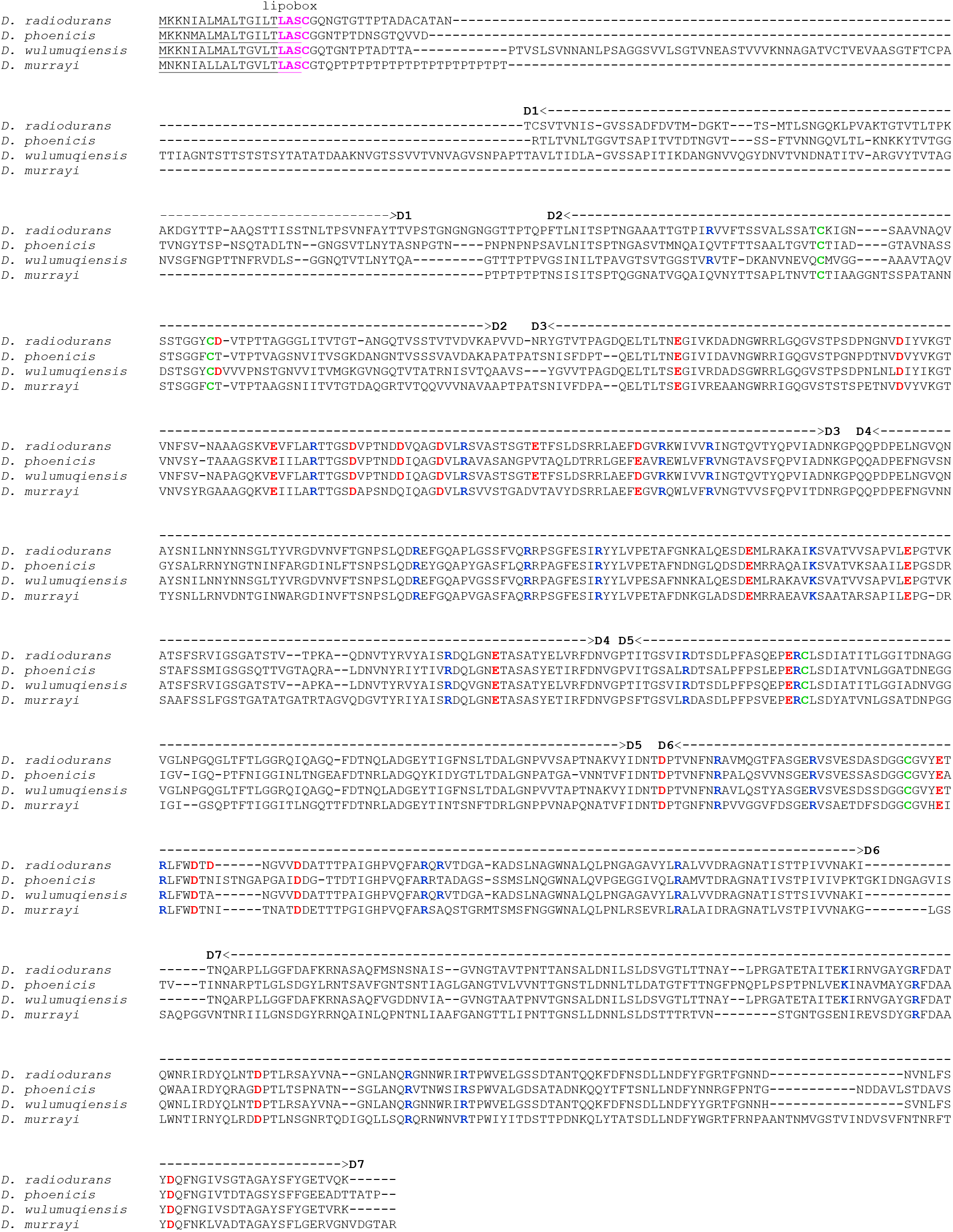
Multiple sequence alignment of *Deinococcal* HPI SLPs. The four-residue-long lipobox motif comprising an invariant cysteine residue is colored pink, whereas positively- and negatively-charged residues observed to form salt bridges in our structure are colored blue and red, respectively. The domain boundaries of the seven Ig-like domains are indicated, and cysteine residues in close proximity in the model are colored green. Accession details for the shown protein sequences are provided in Table S2.

**Figure S3.**
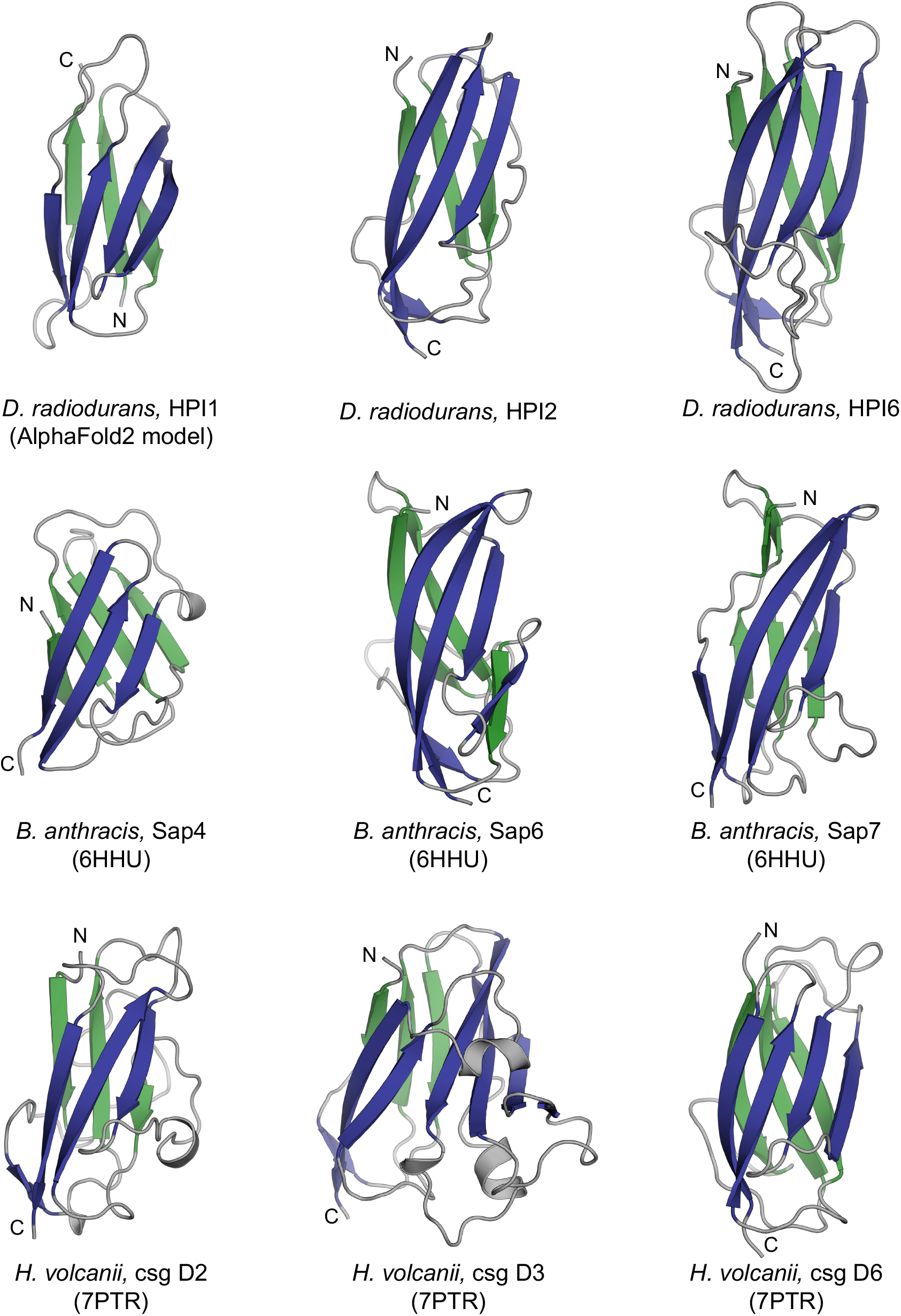
Gallery of Ig-like domain containing SLPs from bacteria and archaea. The structure of *D. radiodurans* HPI1 was predicted using AlphaFold2; the PDB accessions of other shown structures are provided within the rounded brackets. Although quite divergent in their sequences, the Ig-like domains of many prokaryotic SLPs exhibit topologically similar folds.

**Figure S4.**
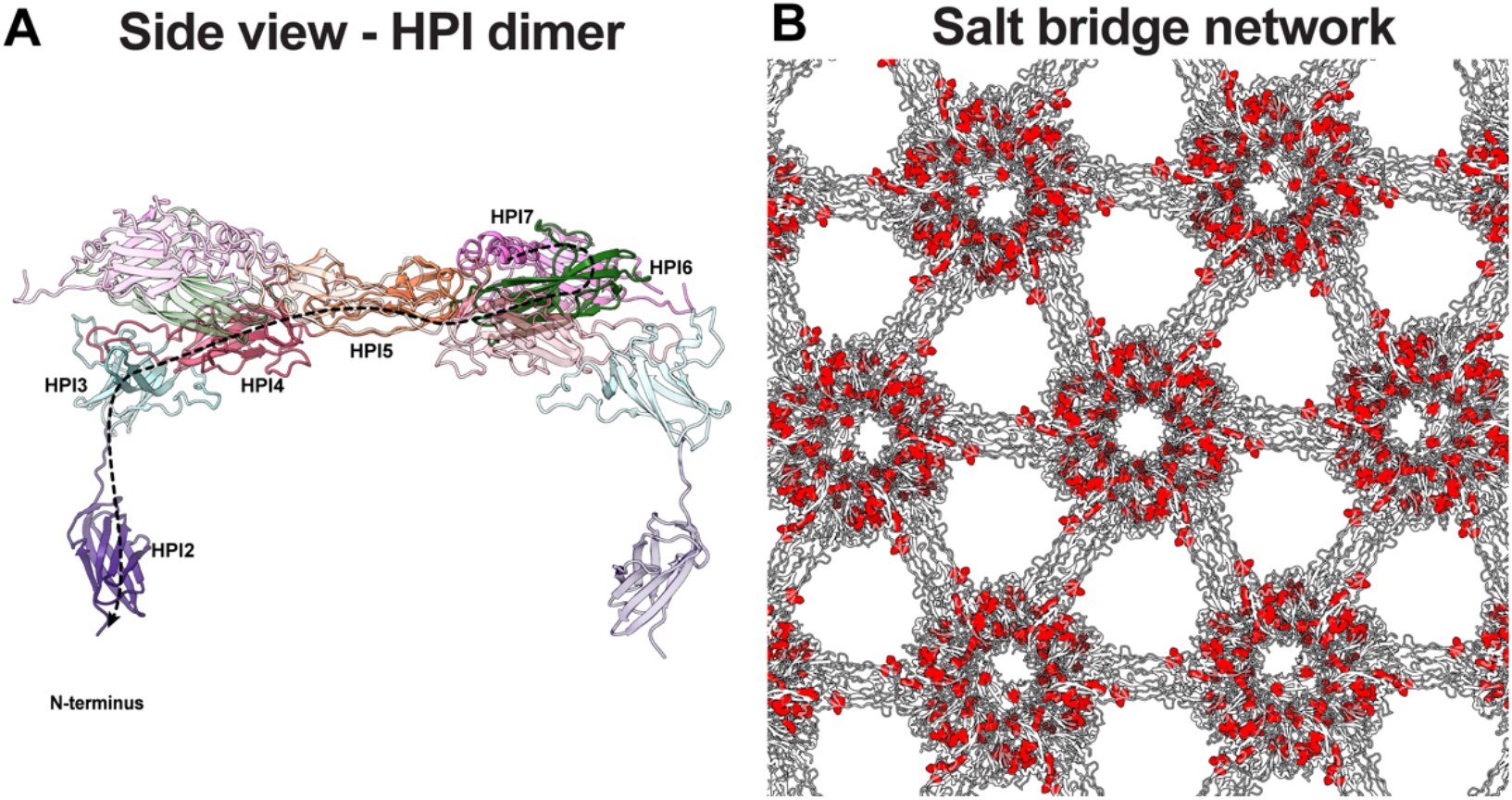
The HPI S-layer lattice arrangement. (A) Side view of the HPI lattice dimer shows how the S-layer is arranged in multiple layers. HPI6-7 form the outermost layer, connected to a lower lattice made up of HPI3-4 through bridging HPI5 domains. HPI2 forms the lowest layer resolved in the map proximal to the cell. (B) The HPI lattice is shown from the top (from outside the cell), with HPI protein residues forming salt bridges colored in red. This extensive salt bridging network stabilizes the lattice.

**Figure S5.**
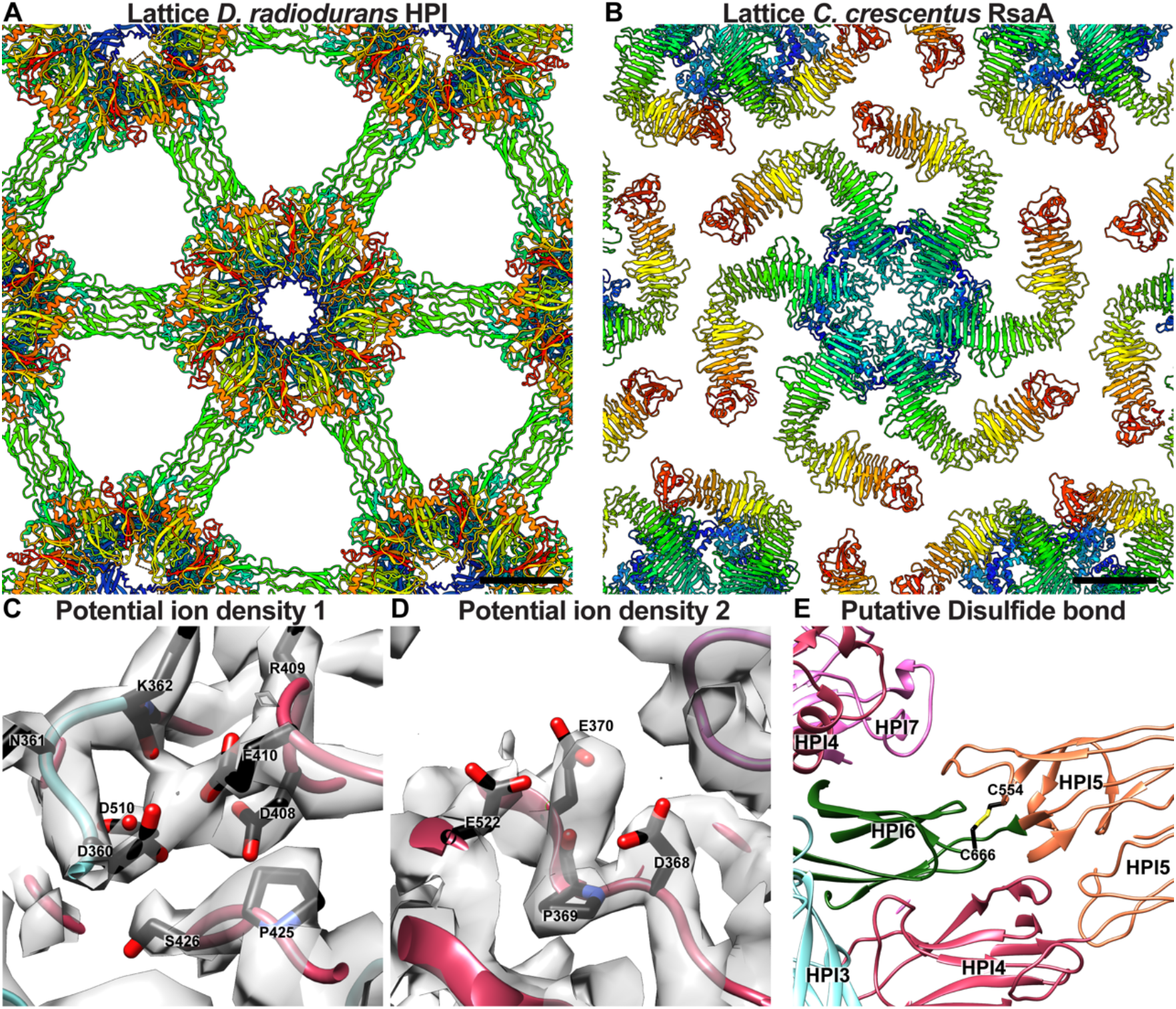
Gaps and putative ion densities observed in the lattice. (A) Top view of the HPI lattice shows several large gaps (pores) in the S-layer (Scale bar 50 Å). Each HPI monomer is colored as rainbow from the N- to the C-terminus. (B) For comparison, a top view of the previously reported outer S-layer lattice from the diderm *C. crescentus* bacterium, where smaller pores are observed. Each SLP monomer (RsaA protein) is colored as rainbow from the N- to the C-terminus (Scale bar 50 Å). (C-D) In several locations, negatively charged residues were bound to unexplained densities, which could be putative cations. (E) Two cysteine residues (C554 and C666) are placed in close proximity in our model, which may further interconnect the lattice with a disulfide bond.

## Supplementary Tables

**Table S1:**
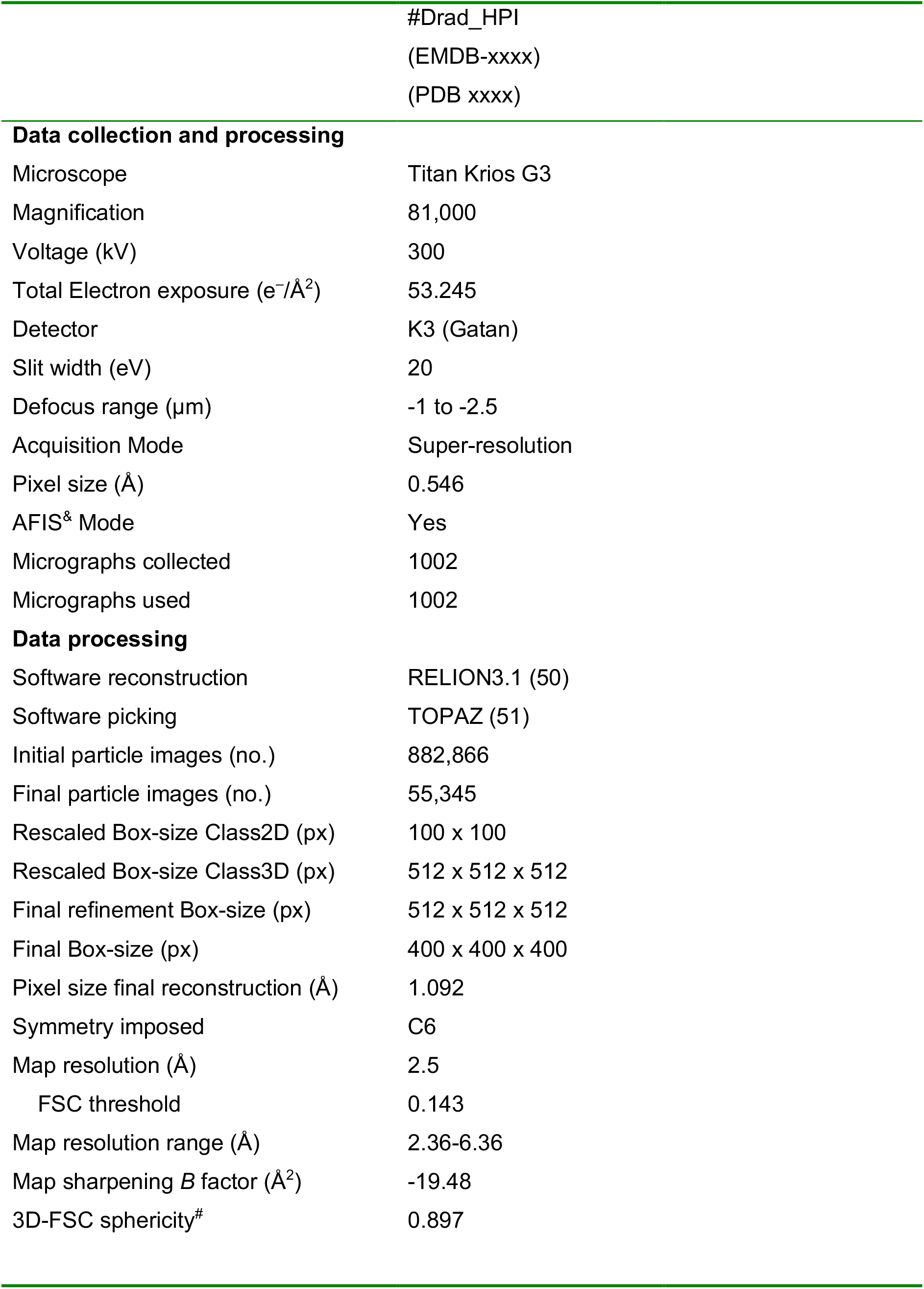
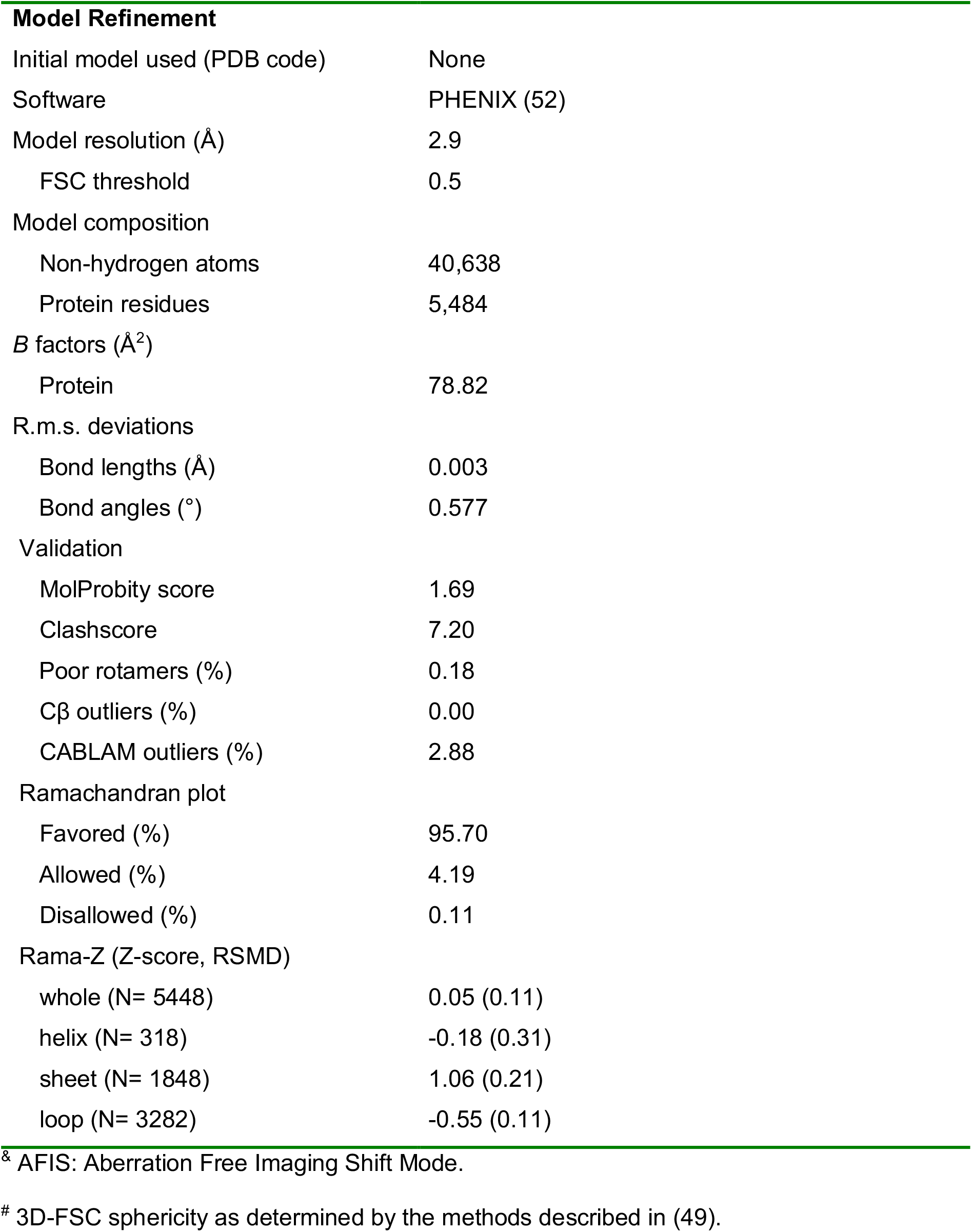
Cryo-EM data collection, refinement and validation statistics.

|  |  |
| --- | --- |
|  | #Drad_HPI<br>(EMDB-xxxx)<br>(PDB xxxx) |
| **Data collection and processing** |  |
| Microscope | Titan Krios G3 |
| Magnification | 81,000 |
| Voltage (kV) | 300 |
| Total Electron exposure (e<sup>-</sup>/Å<sup>2</sup>) | 53.245 |
| Detector | K3 (Gatan) |
| Slit width (eV) | 20 |
| Defocus range (μm) | -1 to -2.5 |
| Acquisition Mode | Super-resolution |
| Pixel size (Å) | 0.546 |
| AFIS<sup>®</sup> Mode | Yes |
| Micrographs collected | 1002 |
| Micrographs used | 1002 |
| **Data processing** |  |
| Software reconstruction | RELION3.1 (50) |
| Software picking | TOPAZ (51) |
| Initial particle images (no.) | 882,866 |
| Final particle images (no.) | 55,345 |
| Rescaled Box-size Class2D (px) | 100 x 100 |
| Rescaled Box-size Class3D (px) | 512 x 512 x 512 |
| Final refinement Box-size (px) | 512 x 512 x 512 |
| Final Box-size (px) | 400 x 400 x 400 |
| Pixel size final reconstruction (Å) | 1.092 |
| Symmetry imposed | C6 |
| Map resolution (Å) | 2.5 |
| FSC threshold | 0.143 |
| Map resolution range (Å) | 2.36-6.36 |
| Map sharpening *B* factor (Å<sup>2</sup>) | -19.48 |
| 3D-FSC sphericity<sup>#</sup> | 0.897 |
| **Model Refinement** |  |
| Initial model used (PDB code) | None |
| Software | PHENIX (52) |
| Model resolution (Å) | 2.9 |
| FSC threshold | 0.5 |
| Model composition |  |
| Non-hydrogen atoms | 40,638 |
| Protein residues | 5,484 |
| *B* factors (Å<sup>2</sup>) |  |
| Protein | 78.82 |
| R.m.s. deviations |  |
| Bond lengths (Å) | 0.003 |
| Bond angles (°) | 0.577 |
| Validation |  |
| MolProbity score | 1.69 |
| Clashscore | 7.20 |
| Poor rotamers (%) | 0.18 |
| Cβ outliers (%) | 0.00 |
| CABLAM outliers (%) | 2.88 |
| Ramachandran plot |  |
| Favored (%) | 95.70 |
| Allowed (%) | 4.19 |
| Disallowed (%) | 0.11 |
| Rama-Z (Z-score, RSMD) |  |
| whole (N= 5448) | 0.05 (0.11) |
| helix (N= 318) | -0.18 (0.31) |
| sheet (N= 1848) | 1.06 (0.21) |
| loop (N= 3282) | -0.55 (0.11) |

<sup>&</sup> AFIS: Aberration Free Imaging Shift Mode.
<sup>#</sup> 3D-FSC sphericity as determined by the methods described in (49).

**Table S2:** NCBI/UniProtKB IDs of prokaryotic SLPs containing Ig-like domain arrays.

| **Organism (protein name)** | **NCBI/UniProtKB ID** |
| --- | --- |
| *Deinococcus radiodurans* (HPI) | P56867 |
| *Deinococcus phoenicis* (HPI) | WP_034356122.1 |
| *Deinococcus wulumuqiensis* (HPI) | WP_162865520.1 |
| *Deinococcus murrayi* (HPI) | WP_027460105.1 |
| *Deinobacterium chartae* (HPI-1) | WP_183984901.1 |
| *Deinobacterium chartae* (HPI-2) | WP_183984905.1 |
| *Deinococcus fonticola* (HPI) | WP_135229161.1 |
| *Thermus thermophilus* (HPI-like) | BDG20071.1 |
| *Bacillus anthracis* (sap) | P49051 |
| *Bacillus anthracis* (EA1) | P94217 |
| *Geobacillus stearothermophilus* (SbsB) | CAA66724 |
| *Geobacillus stearothermophilus* (SbsC) | O68840 |
| *Paenibacillus alvei* (SpaA) | WP_005544766.1 |
| *Acetivibrio thermocellus* | O86999 |
| *Aeromonas salmonicida* | P35823 |
| *Haloferax volcanii* (csg) | P25062 |
| *Halobacterium salinarum* (csg) | P0DME1 |
| *Haloferax mediterranei* (csg-1) | WP_004057339.1 |
| *Haloferax mediterranei* (csg-2) | WP_004059902.1 |
| *Methanospirillum hungatei* | WP_011449226.1 |
| *Methanothermus fervidus* | P27373 |
| *Staphylothermus marinus* (Tetrabrachion) | Q54436 |
| *Aeropyrum pernix* | Q9YEG7 |
| *Sulfolobus islandicus* (SlaA) | F0NHT7 |
| *Sulfolobus islandicus* (SlaB) | F0NHT6 |
| *Metallosphaera sedula* (SlaA) | A4YHQ8 |
| *Metallosphaera sedula* (SlaB) | A4YHQ9 |
| *Nitrososphaera viennensis* (SlaA) | WP_144239589.1 |
| *Nitrososphaera viennensis* (SlaB) | WP_144239588.1 |

## Methods

### Purification of HPI protein

Wild-type HPI protein was purified from *D. radiodurans* bacteria by adapting a previously described protocol (13). Six liters of modified TGY (Tryptone-Glucose-Yeast extract) medium was inoculated 1:50 with a late-log phase culture of *D. radiodurans*, and cells were grown aerobically at 30 °C. Late log phase cells were harvested by centrifugation (5,000 relative centrifugal force (rcf), 4 °C, 30 minutes) and frozen and stored at −80 ºC until further experimentation. The cell pellet from a one-liter culture was carefully resuspended in 80 mL milliQ water, supplemented with 1X cOmplete protease inhibitor cocktail (Roche). The HPI layer was released from the cell surface by adding dropwise a 10% (w/v) SDS stock-solution to the cell suspension up to a final concentration of 2 % (w/v) SDS. The cell suspension was incubated for two hours at room temperature on a rotatory wheel, and solubilized protein was separated from HPI and cell debris by centrifugation (8,000 rcf, 25 °C 20 minutes). The remaining pink pellet was carefully resuspended in 100 mL 2% (w/v) SDS and HPI was separated from stripped cells by centrifugation (3,000 rcf, 25 °C 20 minutes). The remaining supernatant containing HPI was carefully removed from the cellular pellet, and HPI-sheets were subsequently centrifuged (35,000 rcf 25 °C 20 minutes), forming a small opaque pellet at the bottom of the centrifugation tube. The pellet was washed four times by resuspending the pellet in a buffer containing 50 mM HEPES/NaOH pH=7.5, 150 mM NaCl, 5 mM MgCl_2_ and 1 mM CaCl_2_ followed by centrifugation 16,000 rcf 4 °C, 15 minutes. The final pellet was resuspended in the same buffer and the protein concentration was measured by ultraviolet absorption at 280 nm as 4.7 mg/mL. This protein solution was then used for cryo-EM experiments.

### Cryo-EM sample preparation

For cryo-EM grid preparation of purified protein, 2.5 µL of the specimen was applied to a freshly glow discharged Quantifoil R2/2 Cu/Rh 200 mesh grid, adsorbed for 60 s, blotted for 4-5 s and plunge-frozen into liquid ethane in a Vitrobot Mark IV (ThermoFisher), while the blotting chamber was maintained at 100% humidity at 10 ºC. Grids for cryo-FIB milling of *D. radiodurans* cells were prepared as described previously (10). Briefly, *Deinococcus radiodurans* strain BAA-816 (obtained from the ATCC) was grown aerobically in TGY liquid medium (53). Cells were grown for 24 hours at 30 °C prior to harvesting and staining with FM4-64 fluorescent membrane dye (Invitrogen). Four µL of cells were loaded on Finder grids (Electron Microscopy Sciences) and plunge-frozen in a liquid ethane-propane mixture kept at liquid nitrogen temperatures using a Vitrobot Mark IV (Thermo Fisher Scientific). Grids were clipped and stored under liquid nitrogen.

### Cryo-EM and cryo-ET data collection

Cryo-EM single particle data: Single-particle cryo-EM data were collected as described previously (26, 31, 37) on a Titan Krios G3 microscope (ThermoFisher) operating at 300 kV fitted with a Quantum energy filter (slit width 20 eV) and a K3 direct electron detector (Gatan) with a sampling pixel size of 0.546 Å running in counting super-resolution mode. For the HPI sheets S-layer sample used for the hexameric lattice structure, a total of 1002 movies were collected with a dose rate of 4.6 e^-^/pixel/s on the camera level. The sample was subjected to 3.43 s of exposure, during which a total dose of 53.3 e^-^/Å^2^ respectively was applied, and 40 frames were recorded per movie (see Table S1).

Cryo-ET data: cryo-FIB milling of the specimen was performed as described previously (10) using a Zeiss Crossbeam 550 FIB-SEM microscope and generated 200-nm thick lamellae. Subsequently, the EM grids were transferred to a Titan Krios transmission electron microscope (TEM) operating at 300 kV, equipped with a Falcon 3 direct electron detector (Thermo Fisher Scientific) for cryo-ET data collection. Tilt series images were acquired bidirectionally using the SerialEM software (54) at 18,000 x or 22,500x magnification (pixel sizes 4.6 Å or 3.7 Å, respectively) with a defocus of −4 μm, ±60° oscillation, 1º increments with a total final dose of 100 e^−^/Å^2^.

### Cryo-EM single particle analysis

HPI structure from two-dimensional sheets: Cryo-EM data processing was performed as described previously for two-dimensional S-layer sheets (26). Movies were clustered into optics groups based on the XML meta-data of the data-collection software EPU (Thermo Fisher Scientific) using a k-means algorithm implemented in EPU_group_AFIS (https://github.com/DustinMorado/EPU_group_AFIS). Imported movies were motion-corrected, dose-weighted, and Fourier cropped (2x) with MotionCor2 (55) implemented in RELION3.1 (56). Contrast transfer functions (CTFs) of the resulting motion-corrected micrographs were estimated using CTFFIND4 (57). Initially, side views of S-layer sheets were first manually picked along the edge of the lattice using the helical picking tab in RELION while setting the helical rise to 40 Å. Top and tilted views were manually picked at the central hexameric axis. Manually picked particles were extracted in 4x downsampled 100 × 100 boxes and classified using reference-free 2D classification inside RELION3.1. Class averages centered at a hexameric axis were used to automatically pick particles inside RELION3.1. Automatically picked particles were extracted in 4x downsampled 100×100 pixel^2^ boxes and classified using reference-free 2D classification. Particle coordinates belonging to class averages centered at the hexameric axis were used to train TOPAZ (51) in 5x downsampled micrographs with the neural network architecture conv127. For the final reconstruction, particles were picked using TOPAZ and the previously trained neural network above. Additionally, top, bottom, and side views were picked using the reference-based autopicker inside RELION3.1, which TOPAZ did not readily identify. Particles were extracted in 4x downsampled 100×100 pixel^2^ boxes and classified using reference-free 2D classification inside RELION3.1. Particles belonging to class averages centered at the hexameric axis were combined, and particles within 30 Å were removed to prevent duplication after alignment. All resulting particles were then re-extracted in 4x downsampled 100×100 pixel^2^ boxes. All side views and a subset of top and bottom views were used for initial model generation in RELION-3.1. The scaled and lowpass filtered output was then used as a starting model for 3D auto refinement in a 512×512 pixel^2^ box. Per-particle defocus, anisotropy magnification, and higher-order aberrations (50) were refined inside RELION3.1, followed by another round of focused 3D auto refinement and Bayesian particle polishing (50). The final map was obtained from 55,345 particles and post-processed using a soft mask focused on the central hexamer, including the dimeric bridge, yielding a global resolution of 2.5 Å according to the gold standard Fourier shell correlation criterion of 0.143 (58). The two-dimensional sheet-like arrangement led to anisotropy in resolution, with lower resolution perpendicular to the plane as estimated by directional FSCs (49), observed previously by several studies on two-dimensional sheets (59, 60). See also Figure S1 and Table S1 for further details.

### Cryo-ET data analysis

Tilt series alignment using patch tracking and tomogram generation was performed using IMOD (61). Subtomogram averaging was performed using custom scripts written in MATLAB (MathWorks), described previously (35, 62, 63). For the preliminary assignment of angles and initial structure determination, we adopted established methods (32). This workflow allowed us to produce lattice maps from cells. Initial subtomogram averaging maps and output angular angle assignments were then used to refine the subtomogram averages further, resulting in the final maps of the HPI hexamer from cells. Figure panels containing cryo-EM or cryo-ET images were prepared using IMOD and Fiji (64). Lattice maps of S-layers for visual inspection were plotted inside UCSF Chimera (65) with the *PlaceObject* Plugin (66). Supplementary Movie S1 was prepared with UCSF ChimeraX.

### Model building and refinement

HPI hexameric structure from sheets: For model building the original 512×512×512 voxel box was cropped into a 400×400×400 voxel box and the protein backbone of HPI was manually traced as a poly-alanine model through a single HPI subunit using Coot (67). Side chains were assigned at clearly identifiable positions which allowed deduction of the protein sequence register. The atomic model was then placed into the hexameric map as six copies and subjected to several rounds of refinement using refmac5 (68) inside the CCP-EM software suite (69) and PHENIX (52), followed by manually rebuilding in Coot (67). HPI5 formed an extended bridge at the edge of the hexamer, and for that part of the map, special considerations were used for model building. The map in this region was locally sharpened using *servalcat* (70) and this map was additionally rotated using a local two-fold axis to confirm our model building. Model validation was performed in PHENIX and CCP-EM, and data visualization was performed in Chimera, ChimeraX, and PyMOL (71). To analyze lattice interfaces, multiple copies of the hexameric structure were placed in the cryo-EM map prepared with a larger box size.

### Bioinformatic analysis

All sequence similarity searches were performed in the MPI Bioinformatics Toolkit (72) using BLAST (73) and HHpred (74). BLAST searches were performed against the nr_bac database, a version of the NCBI nonredundant protein sequence database (nr) filtered for bacterial sequences, using default settings to identify homologs of HPI in bacteria. The searches were seeded with the protein sequence of *D. radiodurans* HPI (UniProt ID P13126). The domain organization of several obtained matches and many experimentally characterized S-layer proteins (Table S2) were analyzed using HHpred searches in default settings over the PDB70 and ECOD70 databases, versions of the PDB and ECOD databases filtered for a maximum pairwise identity of 70%, and using structural models built with AlphaFold v2.2.0 (27). Signal peptides were predicted using SignalP 6.0 (75).

## Movie Legend

**Movie S1. Atomic structure of the *D. radiodurans* S-layer**.

The cryo-EM map and atomic structure of the *D. radiodurans* S-layer show how the immunoglobulin-like domains of HPI form the lattice. Different views of the S-layer are shown with text annotations.

